# *Streptococcus pneumoniae* rapidly translocates from the nasopharynx through the cribriform plate to invade and inflame the dura

**DOI:** 10.1101/2021.09.28.462246

**Authors:** Teerawit Audshasai, Jonathan A. Coles, Stavros Panagiotou, Shadia Khandaker, Hannah E. Scales, Morten Kjos, Murielle Baltazar, Julie Vignau, James M. Brewer, Aras Kadioglu, Marie Yang

## Abstract

The entry routes and translocation mechanisms of bacterial pathogens into the central nervous system remain obscure. We report here that *Streptococcus pneumoniae* (Sp) or polystyrene microspheres, applied to the nose of a mouse, appeared in the meninges of the dorsal cortex within minutes. Recovery of viable bacteria from dissected tissue and fluorescence microscopy showed that up to at least 72h, Sp and microspheres were predominantly in the outer of the two meninges, the pachymeninx. No Sp were found in blood or cerebrospinal fluid. Evidence that this was not an artifact of the method of administration is that in mice infected by horizontal transmission, Sp were also predominantly in the meninges and absent from blood. Intravital imaging through the skull, and flow cytometry showed recruitment and activation of LysM^+^ cells in the dorsal pachymeninx at 5h and 10h following intranasal infection. Imaging of the cribriform plate suggested that both Sp and microspheres entered through its foramina via an inward flow of fluid connecting the nose to the pachymeninx. Our findings bring further insight into the invasion mechanisms of bacterial pathogens such as Sp into the central nervous system, but are also pertinent to the delivery of drugs to the brain, and the entry of air-borne particles into the cranium.

## Introduction

Infectious diseases affecting the central nervous system (CNS) are among the most devastating illnesses, leading to up to 100% mortality in some cases ^1^. A wide array of infectious agents - of either bacterial, viral, fungal or parasitic origin - can cause infections of the meningeal or parenchymal compartments ^2^. *Streptococcus pneumoniae* (Sp) is a frequent asymptomatic colonizer of the human nasopharynx ^3^, it can spread from there to invade other tissues including the lungs, blood and the cranium, typically when immunity is weakened ^4-6^. Improved understanding of the mechanisms by which neurotropic pathogens, such as Sp, gain access into the CNS would aid the development of more effective preventative or therapeutic strategies.

Intracranial invasion in humans is not examined until clinical symptoms have developed, so there is no direct evidence of the route of initial invasion. In mice, where tissues can be examined at shorter or pre-defined time points, it has been reported that instillation of Sp in the nasal cavity can lead to invasion of cranial tissues in the absence of bacteraemia. Marra & Brigham ^7^ examined homogenized brains of infant rats one hour after nasal instillation and found colony-forming units (CFUs). Rake ^8^, van Ginkel et al., ^9^ and Hatcher et al. ^10^ found that the infection density of brain tissue was greatest in the frontal, olfactory area. This supports the hypothesis that translocation of bacteria from the nasal cavity to the olfactory bulb is through the foramina of the cribriform plate of the ethmoid bone which allow passage of the olfactory nerve bundles ^11,12^ and, indeed, Rake ^8^ states that Sp ‘*appear in the perineural space of the olfactory* ….. *nerve*’. It has been suggested that the infection and inflammation are in the inner layers of the meninges, i.e. the leptomeninx, which is composed of the pia, the subarachnoid space and the arachnoid ^4,13,14^ (Fig. 1A). When Sp is found in the cerebrospinal fluid (CSF), this suggests its presence in the subarachnoid space of the leptomeninx. However, fluid from the outer meningeal layer, the pachymeninx, which contains the collagenous layers that constitute the dura mater ^15,16^, is not normally sampled in the clinic. For instance, the most common presenting feature of pneumococcal meningitis is headache ^17^, and headache involves inflammation of the pachymeninx. The pachymeninx is richly innervated and vascularized, and contains lymph vessels, and, at least over the cortical convexities, is thicker than the leptomeninx ^18-22^. Inoculation of Sp into the ‘subdural space’ of the pachymeninx is an effective route of infection ^15,23,24^. The ‘subdural space’ is now thought to be a virtual space within the pachymeninx situated beneath layers of collagen and above the dural border cells that overlie the arachnoid barrier layer ^15,20,21,25,26^. Here we have distinguished infection of the meninges from that of the brain, and report that, at least at early times, Sp instilled in the nasal cavity reach the dorsal meninges in the pachymeningeal compartment rather than the leptomeninx, and induce an immune response there. We also show that meningeal invasion by Sp without bacteraemia can occur in mice infected through horizontal transmission. To see if translocation from nasopharynx to meninges depended on an active biological feature of Sp, we also looked for (and found) translocation of inert polystyrene microspheres and compared a range of diameters. As well as for microbes, inward translocation from nose to brain is known to occur for stem cells ^27^ and for non-biological particles including neurotherapeutics ^28,29^ and air-borne particulate pollutants ^30^. Our results outline a pathway of entry to the brain that may be common to all of these materials.

**Figure 1.**
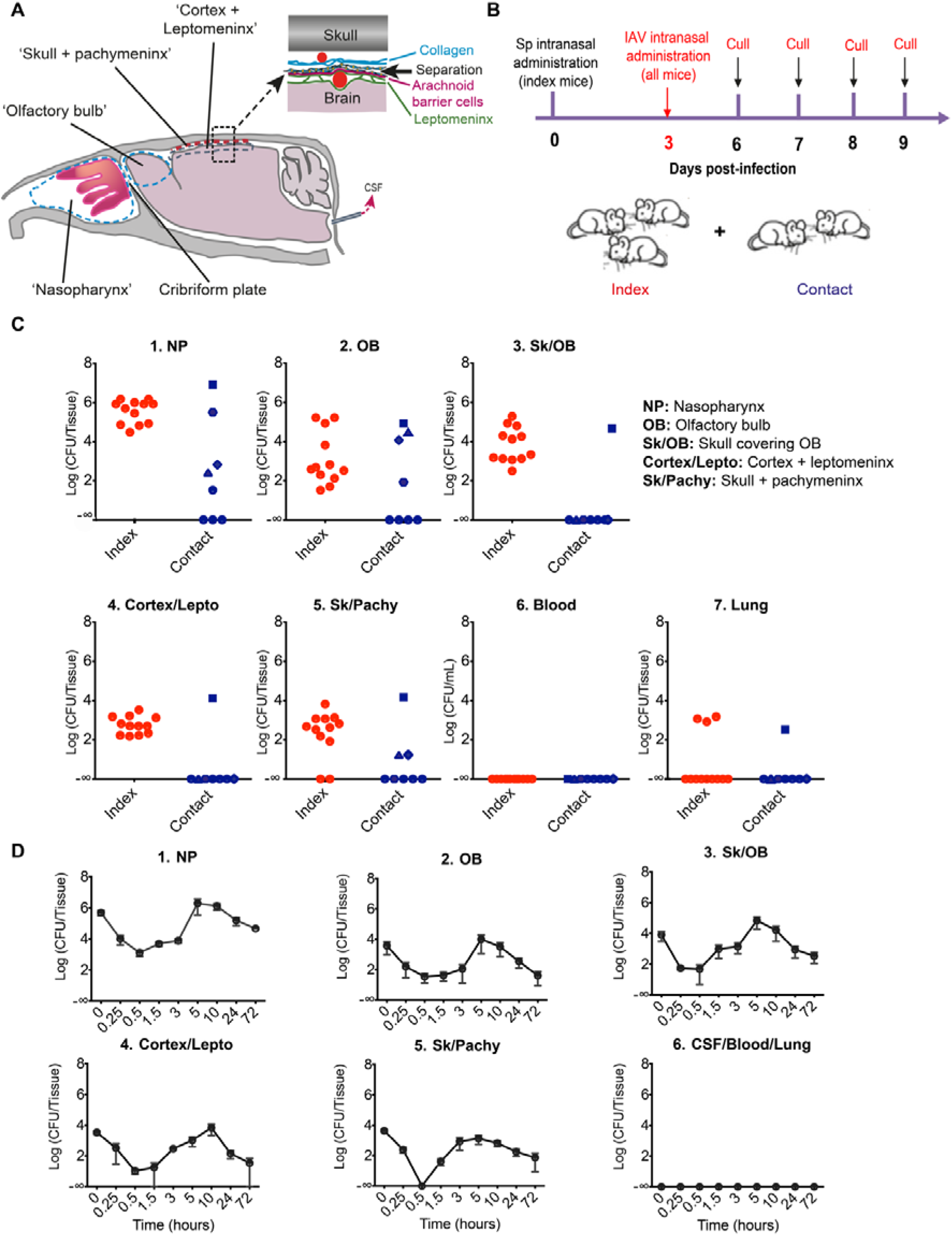
Pneumococci transmitted via horizontal transfer or intranasally instilled reach the meninges, by-passing the blood systemic circulation. (**A**). Schematic representation showing the situation of the tissues investigated. CSF: Cerebrospinal fluid. Top right inset: Schematic magnification of the meninges of the dorsal brain showing the outer layer, the pachymeninx, which contains the collagenous dura mater, the inner layer, the leptomeninx, containing the subarachnoid space, and the underlying brain cortex. Both layers contain blood vessels (red circles). (**B**) Three index mice of a group of five C57/BL6J were infected intranasally with Sp serotype1, strain 217. On day 3, all mice were infected intranasally with Influenza A/HKx31 (H3N2) virus strain. On days 6, 7, 8 and 9, five animals (3 index + 2 contact) per time point were killed with CO_2_ and the number of CFUs per tissue sample counted. **(C**) 1-6: Numbers of CFUs detected per tissue sample from index (n=12) and contact mice (n=8) on days 6, 7, 8 and 9 all confounded. Each dot represents one mouse. (**D**) 1-6: Pneumococci (Serotype 2, strain D39) were intranasally administered and CFUs counted between 0 and 72h in tissue samples as for (C). Data are shown as mean +/- SEM (n=5 mice per time point).

## Materials and Methods

### Ethics statement

All animal experiments were conducted in accordance with the Animals (Scientific Procedures) Act 1986 and Amendment Regulations 2012 (ASPA 2012), and the care and maintenance guidelines of the Universities of Liverpool and Glasgow. All animal protocols were approved by the Local Animal Welfare and Ethics Committees under the UK Home Office Project Licence PB6DE83DA. In line with the 3Rs principle, the number of animals was kept to a minimum and all surgery and intravital imaging were done under terminal anesthaesia.

### Mice

C57/BL6J female mice were obtained from Charles River Laboratories (Kent, UK) at 6-8 weeks old, maintained in an isolator, in a category II animal holding room, and were allowed to acclimatize for at least 7 days before use. LysM^+^ cells were imaged in mice (a kind gift from Professor Sussan Nourshargh, Queen Mary University of London) in which the eGFP gene was knocked into the Lysozyme (Lys) M locus so that myelomonocytic cells were fluorescent, with neutrophils comprising the highest percentage of eGFP^hi^ cells ^31^. CD11c-eYFP mice are described in ^32^.

### Induction of infection

*S*. pneumoniae (Sp) serotype-2, strain D39 (NCTC 7466), were obtained from the National Collection of Type Culture, London, UK. Serotype-1 (sequence type 217) was a clinical isolate obtained from an adult male patient presenting with pneumococcal meningitis, archived at the Malawi-Liverpool-Wellcome Trust Clinical Research Centre, Blantyre, Malawi. The lab-adapted reference strain D39 strain was chosen for its value as a well-characterised benchmark isolate, while Sp serotype 1 was used for its relevance as a high attack rate strain i.e., very short periods of carriage with a high incidence of invasive disease. Both D39 and serotype 1 strains used in this study are known to be viable in blood up to at least 48h when administered intravenously ^33,34^. Bacteria were streaked onto blood agar and grown overnight at 37°C, 5% CO_2_. Sp were identified by presence of a zone of haemolysis round each colony and a zone of inhibition round an optochin disc ^35^. A sweep of colonies was inoculated into brain heart infusion (BHI) broth (Thermofisher) and grown statically overnight at 37°C. The next day 750 µl of overnight growth was subcultured into BHI containing 20% (v/v) fetal calf serum (FCS) and grown statically for 4–6 hr until mid-log phase growth (OD500 0.8), at which point the broth was divided into 500μl aliquots and stored at -80°C in BHI broth with FCS for no more than 1 month until use. Before use, two stock aliquots were thawed at room temperature, serially diluted from 10^−1^ to 10^−6^ and plated onto blood agar plates ^36^ to quantify colony forming units (CFUs).

Mice were anesthetized with 2.5% isofluorane in oxygen and a total of 10 µl of a suspension of Sp in sterile PBS was instilled over 5 - 8 s into the two nares using a micropipette. The mouse was returned to its cage and allowed to recover from anaesthesia. At the chosen time-point it was euthanized in a CO_2_ chamber and tissue dissected as described below. In the horizontal transmission experiments only: three mice of a group of five were infected by Sp, serotype 1, sequence type 217 at a dose of 10^5^ CFU/mouse and returned to their cage. Three days later, all mice were infected intranasally with Influenza virus (IAV) strain A/HKx31 (H3N2) (4x 10^4^ PFU/mouse). IAV is known to promote the rate of pneumococcal transmission (Kono et al., 2016). Tissues were examined for CFUs after a further 3, 4, 5 or 6 days.

In the high-dose intranasal instillation and flow cytometry immunophenotyping study, 10 µl of a suspension containing approximately 10^8^ CFUs of Sp serotype-2 strain D39 was applied in the nostrils and the mice killed with CO_2_ at times comprised between 15 min and 72h. In these sets of experiments, IAV was not administered to any of the mice. After each experimental infection, health checks were performed at least 3 times a day on the infected animals: no visible signs of disease symptoms nor significant changes in motor activity were observed. To determine the viable counts (CFUs) at the shortest time point possible, a group of infected mice were killed by cervical dislocation immediately after recovery from anaesthesia. The interval between the end of the nasal instillation and cardiac arrest was about 2 min 10s. The tissue samples were dissected out 5-7 min later.

### Tissue collection for determination of CFUs

At least 100 µl of blood was taken by cardiac puncture, and 2-5 µl of CSF was collected from the cisterna magna. Four different tissues were taken from each brain and immersed in 1.0 ml of sterile PBS. These were: (1) the dorsal skull, excised with its adhering tissue - the cleavage plane is probably within the inner layers of the pachymeninx ^15^, so some pachymeningeal tissue may have been excluded. This tissue sample is called “skull/ pachymeninx” in Fig. 1. (2) A layer of superficial tissue was sliced off from the dorsal cortex: these samples included the leptomeninx and probably inner layers of the pachymeninx, as well as parenchymal tissue (‘cortex/leptomeninx’). (3) The entire olfactory bulb, and (4) the skull bone overlying the olfactory bulb, with its adherent meningeal tissue, labeled ‘skull/olfactory bulb’. The nasal cavity was exposed by removing the palate, and the nasal septum and associated nasal mucosa were harvested: ‘nasopharynx’. Tissue samples were homogenised using a T10 basic Ultra-Turrax^®^ homogeniser (IKA, Staufen, Germany) running at 30,000 rpm for 6-8 sec at room temperature. 100 µl of the homogenate was transferred to a well on a 96-well plate and ten-fold serial dilutions made in sterile PBS. 60 µl aliquots were spotted on blood agar plates containing 10µg/ml gentamicin. Cerebrospinal fluid (CSF) samples were plated neat. Colonies were counted manually after overnight incubation under anaerobic conditions. To compare densities of CFUs in pachymeningeal tissue scraped from the skull and (cortex + leptomeninx) samples, one volume of lysis buffer (125mM Tris pH 6.8; 5mM EDTA; 1% SDS; 10% glycerol) was added to one volume of undiluted homogenate, and protein content was assayed using a Pierce BCA Protein Assay Kit (Thermofisher) according to the manufacturer’s instructions.

### Fluorescence labeling of *Streptococcus pneumoniae* with CFSE

Sp serotype 2, strain D39 were fluorescence labelled according to a previously described protocol ^37^. After growth to 0.5 OD600 in BHI growth medium at 37°C anaerobically, one ml of the suspension was transferred to a 1.5 ml tube and centrifuged at 4,000g for 5 minutes. The supernatant was discarded and the pellet resuspended in 1 ml of BHI containing 10 µM 5(6)-carboxyfluorescein diacetate N-succinimidyl ester (CFDA-SE, Sigma #21888). The suspension was incubated on a rotating shaker at 37°C and 200 rpm for 45 minutes, in the dark, centrifuged at 12,000g for 3 minutes and washed 3x with room-temperature PBS. The bacteria were resuspended at 10^8^ CFUs/10 µl and stored on ice.

### Post-mortem imaging of the meninges

After the mouse was euthanized, the brain and meninges were perfused through the right cardiac ventricle with either 50 ml PBS or with DiI-glucose solution according to a previously described protocol ^38^, followed by 50 ml of 4% (wt/vol) paraformaldehyde (PFA) solution, at 1.4 ml/min. The lower jaw and the scalp were removed to expose the dorsal and olfactory bulb areas of the skull. Either the underlying soft tissue was left attached and imaged through the skull, or the brain parenchyma and the leptomeninx were removed to leave pachymeningeal tissue which was imaged from the internal face ^39^. The pieces of skull were mounted on a Petri-dish and imaged immediately. Z-stack images were obtained with a Zeiss LSM 880 two-photon microscope with femtosecond excitation at 840 nm with a x10, N.A. 0.3 air or a x20, N.A. 1.0 water immersion objective. CFSE was detected at 500-550 nm, Nile red at 570-620 nm. Image stacks were made at 880 nm excitation wavelength and comprised between 70-240 images with areas up to 425 µm ×425 µm and depths 250-500 µm.

### Baclight^™^ Red-staining and LYVE-1 immunostaining of the skull whole mount

Pneumococci were stained using BacLight™ Red stain (Thermo Fisher), a general cytoplasmic stain ^40^ following the manufacturer’s instructions i.e., 1 μL of a 100 μM DMSO working solution of the BacLight^™^ Red bacterial stain was added to 1 mL of bacterial suspension grown to mid-log phase, followed by 2 washes in PBS (0.1M, pH 7.4) and resuspended in 100μL of PBS. 10μL/mouse of the BacLight^™^ Red-stained Sp suspension was administered intranasally. At 15 min post-administration, mice were sacrificed by CO_2_ asphyxia and perfused with heparin-supplemented PBS solution, followed by 4% paraformaldehyde. The dorsal skull was carefully detached from the brain and trimmed to an area comprising the parietal and frontal bones together with attached meningeal tissue. The resulting tissue was stained using anti-mouse LYVE-1 monoclonal antibody (Thermo Fisher, ALY7-eFluor 450) diluted at 1:200 in PBS, and mounted in a Petri dish for subsequent imaging. The mounted sample was imaged with a Zeiss LSM 880 confocal microscope with excitation set at 561 nm (for Baclight^™^Red) and 405 nm (for LYVE-1). Images were acquired through a x10, N.A. 0.3 air immersion objective. Baclight^™^ Red was detected at 571-664 nm while LYVE-1-eFluor 450 was detected at 416-538 nm.

### Intravital two-photon microscopy through the thinned skull

The microscope and methods were essentially as previously described ^20^. Briefly, the mouse was maintained under isofluorane anaesthesia, adjusted as necessary to suppress the withdrawal reflex, and core temperature was maintained at 37°C with a heating mat. The dorsal skull was exposed and a steel plate with a hole 5 mm in diameter was glued to the skull, usually with its centre about 2 mm caudad to bregma and 2 mm lateral, and held in clamps. In some cases, the mouse was injected through a tail vein with a blood marker such as 70 kD dextran-rhodamine and also furamidine, a nuclear dye which extravasates in the pachymeninx ^41,42^. The skull within the hole in the plate was superfused with Tris buffered saline and thinned with a dental drill. The mouse with attached plumbing was transferred to the stage of an upright two-photon microscope (Zeiss LSM7 MP) controlled by Zen software. The excitation source was a tunable femtosecond laser (Coherent Chameleon Ultra II). This was either used at a wavelength of up to 950 nm or set at up to 880 nm and used to drive an optical parametric oscillator (Coherent) which gave a second beam, typically set at 1140 nm (for mKate). Images were acquired through a x20, N.A. 1.0 water immersion objective (W Plan-Apochromat, Zeiss). Five detector channels were available to separate emission from different fluorophores and from second harmonic generation from bone and collagen. To follow leucocyte movement, Z-stacks about 30 µm deep were collected at intervals of about 30s.

### Image analysis

Two-photon z-stacks and videos were analysed with Imaris 9.5 (Bitplane) and Fiji (NIH Image) software packages. To separate neutrophils from other, less bright, cells in LysM+-eGFP mice, each movie was normalized to the same mean brightness and contrast was set manually against images obtained from an uninfected mouse. Cells were further selected for XY diameters 12 μm or greater and identified as neutrophils. The approximate mean speeds were calculated from their positions in sequential 30 µm z-stacks obtained at the minimum repetition interval, typically 30s. The number of LysM^+^ within each z-stack were quantified using the ‘spots’ function. Values were then converted to the number of cells per mm^2^ according to the size of the imaging area. In order to enhance signal inside the region of interest (below the skull), the surface rendering of the skull as visualised by the SHG was generated and any signal found above this generated surface (outside the skull), was set to 0. To measure the distances from the skull to fluorescent Sp (or microspheres) in the 3D reconstructions, the distance measurement function of Imaris was used to calculate the shortest distance from the center of the positive signal to the surface rendering of the skull.

### Flow cytometric analysis

Groups of C57BL/6J female mice (n = 5/time point) were infected with *S. pneumoniae* D39, and euthanized at post-infection times ranging from 1h to 18h. Pachymeningeal tissue was scraped from the calvaria, gently crushed and passed through a cell strainer to produce a single cell suspension in Dulbecco’s phosphate-buffered saline (Thermofisher). Cells were counted and stained with anti-mouse antibodies to CD45 (clone 30-F11, BD Biosciences), CD4 (clone RM4-5, Biolegend), CD11b (clone M1/70, eBioscience), CD11c (clone N418, eBioscience) and LySM D1 (clone G3, Santa Cruz Biotechnology,) or Ly6G (clone 1A8, Biolegend), in the presence of anti-CD16/32 Fc-receptors block (BD Biosciences). Events were acquired using a FACS Canto II (BD Biosciences) flow cytometer (Supplementary Fig. S4).

### Intranasal administration of microspheres

Fluorescent polystyrene microspheres of three nominal diameters were used: yellow-green (505/515 nm, Thermofisher F13081, 4 × 10^10^ microspheres/ml) and Nile Red (Thermofisher F8819, 4 × 10^10^ microspheres/ml) nominally 1 µm with carboxylate modified surface, Nile Red nominally 5 µm (5-7.9 µm) (Spherotech, FP-6056-2, 1.5 × 10^8^/ml) and Nile Red nominally 10 µm (10-14 µm) (Spherotech, FH-10056-2, 10^7^ microspheres/ml) with unmodified surfaces. For imaging and flow cytometry on meningeal tissue, suspensions containing approximately 10^7^ microspheres per ml were prepared and a volume of 10µl (containing 10^5^ microspheres) was applied intranasally to each of five mice for each diameter and the mice were culled by CO_2_ asphyxia 30 min later. In experiments designed label the leptomeninx, microspheres were injected in the cisterna magna using a 34G 10µl microsyringe (Hamilton). All of these experiments involving microspheres were conducted without any prior or subsequent co-infection with either Sp or IAV.

Two-photon imaging was done through the skull and into the meninges in the areas of the olfactory bulb and the dorsal cortex, or from the intracranial face of the skull with its attached pachymeningeal tissue. Excitation was at 840 nm which produced two-photon excitation of the fluorophores and second harmonic generation (SHG, in blue). For flow cytometry, pachymeningeal tissue scraped from the dorsal was collected in PBS and passed through a cell strainer. A BD Canto II flow cytometer detected the YFP- and Nile red-labelled microspheres using the FITC and PerCP Cy5.5 channels, respectively.

### Statistical analysis

For comparison of multiple groups, the statistical significance of endpoints was evaluated by one-way ANOVA followed by Tukey’s multiple comparisons *post hoc* test. For comparison of two groups, the unpaired two-tailed Student’s *t* test was used. Data are presented as means ± SEM in bar graphs. Statistical significance was reported as *, P < 0.05), **, P < 0.01), ***, P < 0.001; ****, P < 0.0001). All statistical analyses were performed with Prism software (version 8.0, GraphPad Software).

## Results

### Translocation of pneumococci following horizontal transmission

To examine the dissemination of Sp in mice through horizontal transmission, infected (index) mice and uninfected (contact) mice were housed together. A suspension of Sp, serotype-1, sequence type 217, was applied to the nares of three index mice (adult C57/BL6J) in a cage of five (Fig. 1B). Three days later, influenza A virus (IAV) was applied to the nares of all five mice. It is known that, at least in infant mice, the inflammation caused by IAV facilitates dissemination of Sp away from the nasopharynx to other tissues ^43^ and increases shedding via nasal secretion ^43-46^. Four different groups of mice i.e., 5 mice per time point i.e., 3 index and 2 contact mice, were euthanized on days 6 - 9 (Fig. 1B) and blood and tissues analyzed for colony forming units (CFUs). The pooled results for the twelve index mice and eight contact mice are shown in Fig. 1C. All the index mice showed colonization of the nasopharynx, and so did five of the eight contact mice, a transmission rate of 62.5% (Fig. 1C1). Dissemination in the cranium was examined in four tissue samples for each mouse: the dorsal skull bone and the soft tissue that remained attached to it when it was separated from the brain, the superficial cortex underlying this with its attached meningeal tissue, the frontal skull overlying the olfactory bulb and attached tissue, and the olfactory bulb with attached meningeal tissue (Fig. 1A). Pneumococci were found, in one or more mice, in all of these four tissue selections (Fig. 1C 2-5). Since mechanical separation of skull from brain appears to split the meninges at the inner layers of the pachymeninx ^15,47^, the “pachymeninx” samples probably included most of the pachymeninx while the meninges attached to the brain tissue were the leptomeninx, probably contaminated with some pachymeningeal tissue. Viable pneumococci were found in the lung tissue of one contact mouse as well as three index mice (Fig.1C7), but in neither index nor contact mice were CFUs obtained from blood (Fig. 1C6). In other experiments, in which IAV was not administered, horizontal transmission still occurred but at a lower transmission rate and no viable pneumococci were found in the meninges (Supplementary Fig. S1). In transmission experiments similar to those presented here but using infant mice and bioluminescent Sp, Diavatopoulos et al. ^43^ detected luminescence *in vivo* from the lungs but not from within the cranium. A possible explanation of the difference is that since the meninges are very thin, the total numbers of bacteria they contain are small relative to bulky tissues such as the lungs, and therefore would be difficult to detect by *in vivo* bioluminescence. To our knowledge, this is the first report that upon horizontal transmission, pneumococci can translocate from the nasopharynx to the cranium of mice without bacteraemia.

### Translocation of pneumococci following direct nasal application

The horizontal transmission model came with the challenging task of determining the precise timing and anatomical route of infection. Hence in order to better characterize the nasopharynx-to-meninges translocation and its time course, we had recourse to intranasal instillation, a widely used procedure for studying the entry of pathogens into the central nervous system. We infected mice at a defined time by applying a suspension of Sp to the nares (without co-infection with IAV). The same tissues as those analyzed in the horizontal transmission model were collected over a period of up to 72h (Fig 1D 1-6). All the mice showed localized infections, in, at least, the nasopharynx, but at no time point was bacteraemia detected, nor were CFUs found in cerebrospinal fluid (CSF) samples recovered from the cisterna magna, or in lung tissue (Fig. 1D6). At the earliest time point, when the mouse was killed about 2 min after nasal application and the tissue dissected immediately, CFUs were found not only in the nasopharynx (Fig. 1D1) but also in the olfactory bulb and attached meninges (Fig.1D2). Since the translocation from nasopharynx to cranium is so fast (minutes), it is unlikely that it involves damage to cells of the nasopharynx. Apart from the fact that there was no detectable release of pneumococci to the blood, a number of studies have indeed shown that pneumococci cause detectable damage to cells only after several hours of exposure ^33,48-55^. Surprisingly, CFUs were also recovered at this earliest time point from the more remote tissue associated with the dorsal skull and cortex. In all these tissues, the numbers of CFUs then fell by about two log units to reach a minimum at the 30 min time point, even, in the skull/pachymeninx sample, becoming undetectable. The numbers of CFUs then increased with a doubling time of less than 20 min to reach a peak at 5-10h, before falling again. CFU counts then declined gradually over time, but persisted up to 14 days post-infection, including in the brain and leptomeninx (Supplementary Fig. S2). This time course (Fig. 1D) is different from that reported by Dommaschk et al. ^56^, who found a monotonic decrease over 28 days. However, in that study, the initial measurement was not done until 1 h after nasal instillation and they used a different strain of Sp (serotype 3, A661).

### After translocating from the nasopharynx to the dorsal meninges, pneumococci are predominantly found in the pachymeninx but outside lymph vessels

To quantify the density of viable Sp in the pachymeningeal tissue adhering to the skull, the soft tissue was scraped from the skull and the number of CFUs expressed per mg protein content. This was compared with the density in the superficial cortex and attached leptomeninx. At 10h post-infection, in the five of eight mice that presented CFUs, the mean number in the pachymeningeal tissue was 276 times (SD=162, p = 0.019) that found in the cortex samples (Fig. 2A). This shows that Sp were far more concentrated in the pachymeninx than in the superficial cortex and attached meninges. Indeed, if the separation occurred in the inner layers of the pachymeninx, it is possible that all the intracranial CFUs in these mice were in the pachymeninx, the CFUs in the ‘cortex’ samples coming from contaminating remnants of the inner pachymeninx.

**Figure 2.**
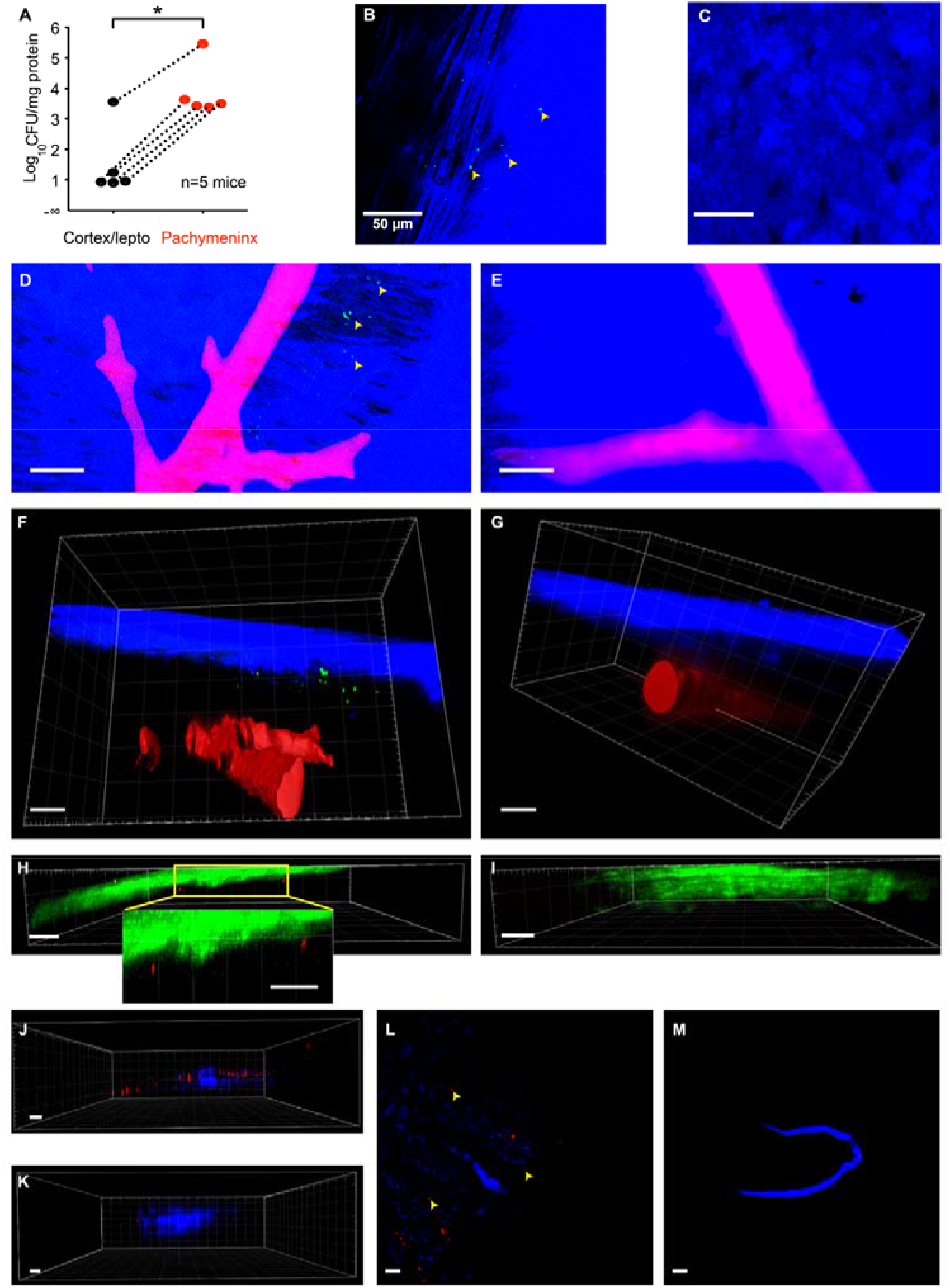
Intranasally administered pneumococci are rapidly and predominantly found in the pachymeningeal compartment of the dorsal meninges. (**A**) Pneumococci were applied to the nose and the mice euthanized 10h later. The skull and brain were separated and CFUs were counted for tissue from the superficial cortex + attached tissue, and for tissue scraped from the skull (‘pachymeninx’). The numbers of CFUs were counted and expressed relative to the weight of protein per tissue sample. In the infected mice, the density of CFUs was much higher in the tissue scraped from the skull. (**B**) CFSE-labelled Sp 90 min post-infection in tissue adhering to the skull after removal of the brain and imaged from the intracranial face (the ‘amaguri’ preparation of Toriumi et al., (2011). Excitation was by laser at 840 nm which produced two-photon excitation of CFSE (green), and Second Harmonic Generation SHG (blue) from collagen and skull bone. Image representative of n=3 mice. Maximum intensity Z-projection, z=171 µm. (**C**) Under identical imaging conditions to (B), no green particles were detected in an uninfected naive mouse. Maximum intensity Z-projection, z=178 µm. (**D, F**) CFSE-labelled *S. pneumoniae* were instilled in the nose of a mouse. Thirty minutes later, the mouse was killed with CO_2_ and perfused with DiI to label blood vessels (Li et al., 2008). Imaging was done through the skull and into the meninges. CFSE-labelled Sp (green) are seen in a z-projection 444µm deep (D). In a 3D projection Sp are seen close to the skull (blue:SHG) and above large blood vessels (red), but are absent from deeper layers (F). (**E, G**) Under identical imaging conditions to (F), no green particles were detected in an uninfected mouse. Z-stack (E) and 3D representation (G). Scale bar: 50µm. (**H**) *S. pneumoniae* expressing mKate were instilled in the nose. After 3.7h, the dorsal meninges and underlying brain were imaged *in vivo* through the skull with excitation at 1140 nm. The SHG from skull bone and collagen is green and emission from mKate is red. A representative XZ section including two groups of pneumococci is shown. (**I**) Similar red signals were not seen in uninfected mice under the same imaging conditions as (H). (**J, K)** *S. pneumoniae* stained with BacLight™Red were instilled in the nose. At 15 min post-administration, mice were perfused transcardially with PBS followed by fixing solution (4% PFA). Dorsal skull mounts were stained with anti-LYVE1 antibody and imaged on the skull bone-oriented surface with excitation at 561 nm and 405 nm. A representative XZ projection of the skull whole mount is shown for the Sp-infected (J) and uninfected (K) mouse. (**L, M)** Maximum intensity Z- projection of the images shown in (J) and (K), respectively z=16.75 µm and z=340.37 µm.

To obtain further information on the location of intracranial Sp, we used two-photon microscopy and fluorescent Sp. In the first method, Sp were labeled in culture by uptake of carboxyfluorescein succinimidyl ester (CFSE) (Supplementary Fig. S3A) and applied to the nose. Thirty minutes later, the mouse was killed with CO_2_, the brain removed, and a piece of dorsal skull bone with adherent tissue imaged from the intracranial side ^39^. With femtosecond excitation at 840 nm, sparse particles emitting green fluorescence were visible (Fig. 2 B); these were in about the same plane as the collagen fibres of the dura made visible by second harmonic generation (SHG) ^41^. Green particles were not seen in uninfected (i.e. naïve) mice (Fig. 2 C). To see if the distribution of Sp extended deeper under the skull than the pachymeninx, CFSE-labeled Sp were also imaged through the skull into the intact meninges and superficial parenchyma. To provide anatomical markers, blood vessels were labeled by intravenous infusion of the carbocyanine dye DiI ^38^. Again, fluorescent particles were observed in tissue from infected mice (Fig. 2 D) but not uninfected ones (Fig. 2 E). In 3D reconstructions, it was evident that the green particles in infected mice were close to the skull and above the pial blood vessels (Fig. 2 F,G). Although endogenous fluorescent particles can often be seen with two-photon microscopy, their detection requires a higher excitation intensity and detector sensitivity than those used here. Nevertheless, to check that the CSFE-labeled Sp were not confused with endogenous fluorescent objects, we also used Sp with very different excitation and emission spectra. Mice were infected with Sp expressing the red fluorescent protein mKate2 ^57^. Ten hours later, the dorsal meninges and superficial cortex were imaged *in vivo* through the skull using two-photon excitation at 1140 nm. Red particles were observed close below the green SHG of the skull of infected mice (Fig. 2 H) but not in uninfected mice (Fig. 2 I).

To examine the location of pneumococci in relation to dural lymph vessels, we have used yet another label to stain pneumococci (BacLight™ Red, ThermoFisher ^40,58^) which appear as dots in the meninges of infected mice (Fig 2J-K, Supplementary Fig. S3B), while none were found in uninfected mice (Fig. 2L,M). Our results clearly show that pneumococci that reach the pachymeninx from the nasopharynx are located outside, not inside, LYVE-1+ structures. Since the meningeal lymph vessels are in the pachymeninx ^59-61^, the presence of a LYVE-1 signal close to the BacLight™Red-labelled pneumococci is further evidence that Sp are in the pachymeninx.

We next sought to determine the distance from the skull of the fluorescent signals. The mean measured distance of Sp (CFSE- and mKate2-labelled) was 18.2 µm, SD 13.6 µm, N = 120 particles measured in 4 z-stacks and 4 mice (Fig. 3D, Sp black circles). Although the large blood vessels in the pachymeninx and in the leptomeninx make the thicknesses of both layers very variable ^20^, the distribution of depths suggests that most pneumococci were in the pachymeninx, and certainly not within the brain parenchyma. As expected, Sp labelled by uptake of CFSE lost fluorescence as the dye was diluted in successive generations ^62^. On some occasions, expression of mKate2 was also lost but we found that robustly fluorescent polystyrene microspheres of diameter 1 µm were also transported from the nasopharynx to the meninges so we used these as a tentative surrogate for Sp. It is known that Sp and molecules infused in the mouse cisterna magna are carried by CSF to spaces in the leptomeninx ^20,63,64^. Hence to make another test of whether particles entering from the nasopharynx were arriving in this space or in the pachymeninx, we applied green fluorescent microspheres to the nose and infused red fluorescent microspheres into the cisterna magna of the same mouse. 30 min later the mouse was euthanized and the meninges and cortex were examined through the skull with two-photon microscopy (Fig. 3A-C). The microspheres administered intranasally were found at a mean distance from the skull of 24.2 µm, SD 13.6 µm, N = 90 particles measured in 4 z-stacks in 4 mice, which is not significantly different from that of the bacteria (Fig. 3D). The mean depth of those infused in the cisterna magna was 81.0 µm, SD 15.9 µm, N =6 particles measured in 3 z-stacks in 3 mice, which is more than three times greater (Fig. 3D). Altogether, the various techniques we used concur in showing rapid translocation from the nasopharynx to the pachymeninx of the dorsal meninges.

**Figure 3.**
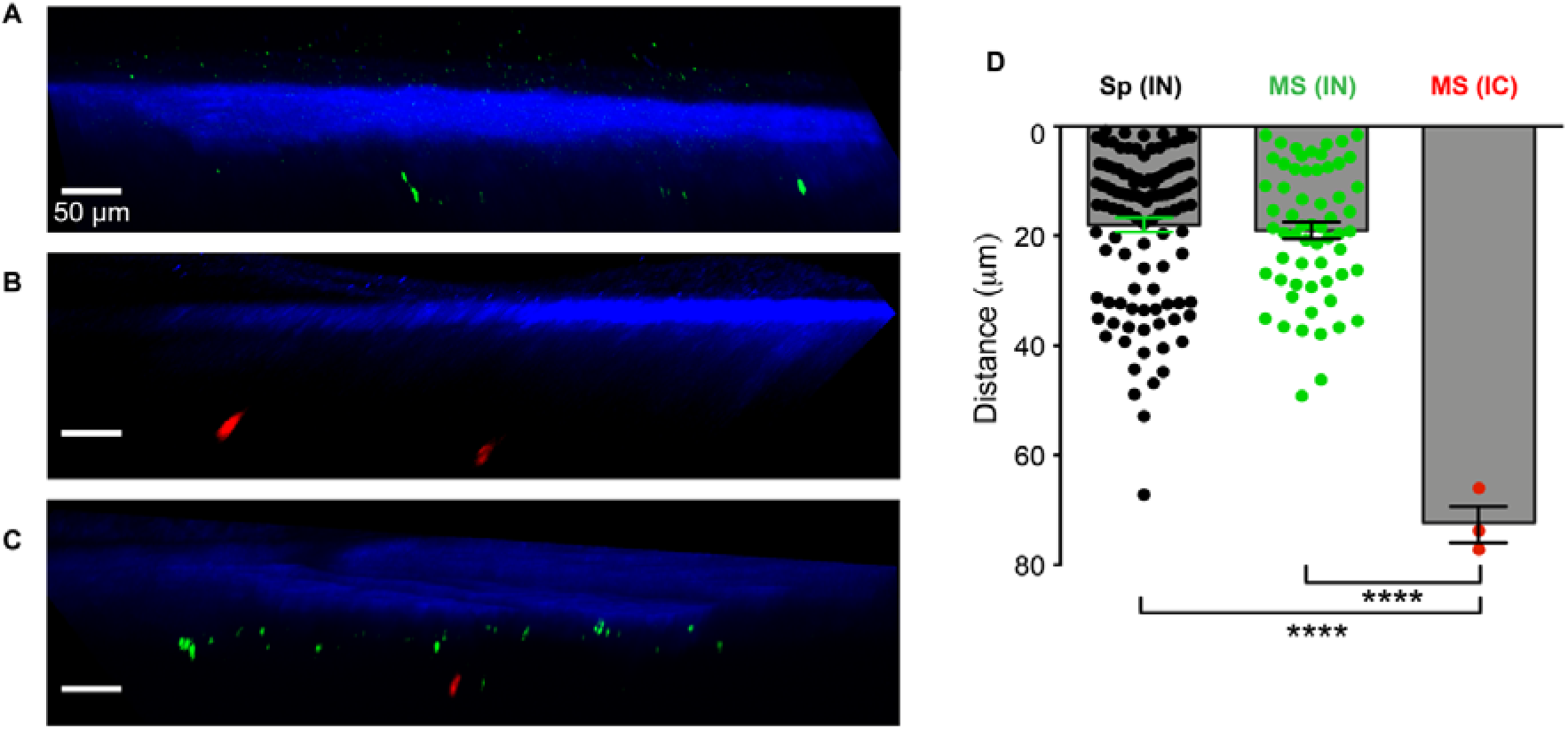
Microspheres administered intranasally were found in a layer close to the skull. (**A-C**) One-micron diameter green fluorescent microspheres were applied to the nose of a mouse (**A**) and 2µm Nile Red-labelled microspheres were injected in the cisterna magna of another mouse (**B**). A third mouse was subjected to both procedures, i.e., intranasal administration of 1µm green fluorescent microspheres followed immediately by intracisternal injection of 2µm Nile Red-labelled microspheres (**C**). In each case, 30 min after the infection, the mouse was euthanized, and the meninges and cortex were examined through the skull with two-photon microscopy. Microspheres administered intranasally were only found in a layer close to the skull (A,C) while those infused in the cisterna magna were deeper (B,C). Scale bar: 50µm. (**D**) The distances from the skull of all fluorescent Sp (black circles) measured in the 3D reconstructions measured post mortem as for Figure 2, panels D-I. The distances from the skull of Sp and fluorescent microspheres measured in the 3D reconstructions measured post-mortem as for Figure 3, panels A-C, upon intranasal instillation of Sp (black circles, n=120 signals, imaging of 4 mice), or upon intranasal (green circles, n=90 signals, imaging of 4 mice) or intracisternal (red circles, n=6 signals, imaging of 3 mice) administration of microspheres. ****p<0.0001, IN: Intranasal, IC: Intracisternal, MS: Microspheres, Sp: *Streptococcus pneumoniae*.

### Nasal administration of pneumococci causes recruitment of innate immune cells to the dorsal pachymeninx

In extracranial tissues such as lung and spleen, Sp and neutrophils interact vigorously ^65-68^. To see how neutrophils reacted to arrival of Sp in the dorsal meninges and superficial cortex, we imaged them *in vivo* by intravital two-photon microscopy through the thinned skull of mice expressing eGFP under control of the LysM promoter (Fig. 4 A-D). In addition to neutrophils, LysM is expressed in other cells of the myelomonocytic lineage, but in *LysM*^*GFP/GFP*^ mice, neutrophils are the brightest ^31^ and can be distinguished from macrophages ^66,69^. In agreement with others, we found very few LysM+ cells in uninfected mice ^69-72^ (Fig. 4D,E), confirming that the skull-thinning and two-photon imaging did not recruit myelomonocytic cells to the meninges within the duration of the experiment ^71^. After nasal infection with pneumococci, the number of LysM+ cells in the dorsal meninges was increased at 5h and 10h (Fig. 4 E). Nearly all of them were in a layer close under the skull, in the plane of smaller vessels typical of the pachymeninx ^73,74^ (Fig. 4 C, D), and above the pia and parenchyma (Fig. 4 A, B) where the microglia are present ^75^. Analysis of videos (Supplementary Videos 1, 2) showed that motile LysM+ cells moved at progressively higher mean speeds as infection progressed (Fig. 4 F) (see also z-projections of tracks in Fig. 4 C and D). The mean speed of motile LysM+ cells at 10h post-infection was 10.4 ± 0.4 µm/min, which is close to the 9.7 µm/min found by Kreisel et al. ^66^ for neutrophils in mouse lung. Many LysM^+^ cells in the pachymeninx followed generally directed trajectories, some along the outsides of blood vessels (Fig. 4 C, Supplementary Video 2) rather than making random walks ^66^. The averaged x, y, and z components of the velocities were not significantly different from zero, i.e., no global drift was detected.

**Figure 4.**
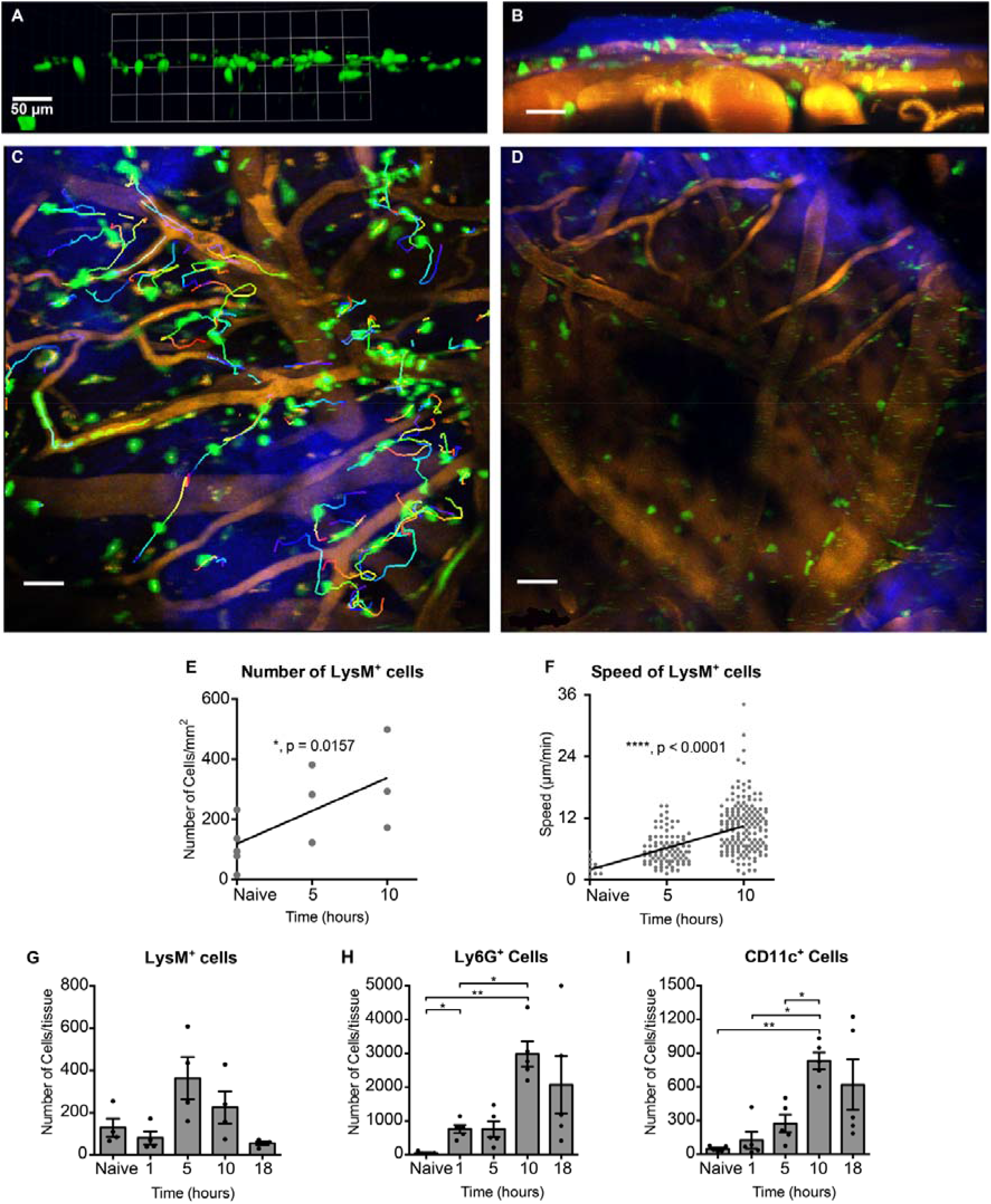
Intranasal infection by *S. pneumoniae* leads to transient recruitment and activation of LysM^+^ in the calvarial pachymeninx. **(A, B**). In vivo two-photon imaging shows that nearly all intracranial LysM^+^ cells are in the meninges. (**A)**. Horizontal view of a 3D reconstruction from a Z-stack of an uninfected mouse showing only LysM^+^GFP cells, which lie in a shallow layer. (**B)**. A different view of the 3D image in (**A**) showing, in addition to the LysM^+^GFP cells (in green), the skull bone (in blue: SHG), nuclei of the pachymeninx (blue from intravenous injection of furamidine) and blood vessels (shown orange-yellow, labeled with rhodamine). Excitation at 840 nm. **(C)** Tracks of LysM^+^ cells in the meninges of a mouse imaged at 10h after intranasal administration of pneumococci. Z-projection of Z stacks 23 µm deep, time series for 15 min. (**D**) Tracks of LysM+ cells in the meninges of an uninfected mouse under the same imaging conditions as (C) z-projection 30 µm deep, time series for 32 min. (**E**) Numbers of LysM^+^ GFP cells per unit area of the meninges counted in *in vivo* images. Each point was obtained from one Z stack. The linear regression line has a slope greater than one with P = 0.016. (**F)** Mean track speeds of mobile GFP^+^ cells in the same imaging conditions as **(E)**. (**G-I**). Flow cytometry of cells from tissue scraped from the calvarial skull. Cells selected as CD45+, CD4+ and CD11b+ were further sorted into LysM+ (G), Ly6G+ (H) or CD11c+ (I) cells. Each dot represents one mouse, error bars are SEMs.

To determine the changes in numbers of LysM^+^ cells in the dorsal pachymeninx over a wider range of times (0 to 18h) we used flow cytometry of tissue scraped from the dorsal skull of C57BL/6 wild-type non-reporter mice. In addition to LysM (Fig. 4G), other gating was used to select cells expressing Ly6G, an integrin-binding protein strongly expressed only on neutrophils ^76,77^, although detectable on eosinophils ^78^, and also CD11c, a marker of dendritic cells (DC) and macrophages ^32,79^ (Fig. 4 H, I). In agreement with the intravital imaging (Fig. 4 E), the number of LysM+ increased over about 5-10h then tended to decrease (Fig. 4 G and H). The time course of Ly6G expression appears to be delayed compared to that of LysM. Since the expression of the *lysM* gene is driven differently from that of *ly6G*, changes in their relative quantities might be expected as the neutrophil population responds to the presence of Sp ^31,80^. As well as LysM^+^ cells, the number of CD11c^+^ cells also increased after nasal infection (Fig. 4I). In videos of the dorsal meninges of infected CD11c-eYFP reporter mice (Supplementary Video 3) nearly all the YFP^+^ cells displayed a rapid extension and retraction of dendrites, suggesting that they, and therefore most of the CD11c^+^ cells of Fig. 4I, were dendritic cells. The number of CD11c+ cells in the dorsal pachymeninx increased some ten-fold, to a peak at about 10h (Fig. 4I). This increase is much earlier than those reported in the nasopharynx and nose-associated lymph nodes, which were insignificant until 3 weeks after nasal infection with Sp ^56^. It is also much faster than the increase in pachymeningeal DCs caused by trypanosomiasis, which occurs between 5 and 10 days after infection ^41^.

### The speed of translocation from nasopharynx to meninges is size dependent

Unlike pneumococci, chemically inert microspheres are not susceptible to destruction by the host’s defenses or by fixation of the tissue, nor can they multiply, hence tracking is simplified. In addition, the brightness and stability of the microsphere fluorescence gives more confidence that the signals detected by microscopy were not artifacts. In Fig. 3 A,C, it was shown that fluorescent polystyrene microspheres with a diameter of 1 µm, close to that of Sp, reached the dorsal pachymeninx from the nasal cavity in under 30min. Since they reached the same destination as Sp and with similar rapidity, we hypothesize that they may have been transported in the same way. To obtain clues to the mechanism of translocation, we asked if it could support microspheres of diameter greater than 1µm. We therefore tested and compared the translocation of microspheres of diameters 1, 5 and 10 µm. At 30 min after nasal administration, microscopic observation of microspheres in the meninges overlying the olfactory bulb suggested abundance in the order 1µm<5µm<10 µm (Fig. 5 A) while, in contrast, in the dorsal meninges the order of abundance was 1µm>5µm>10µm (Fig. 5 B). These distributions were quantified by flow cytometry on pachymeningeal tissue scraped from the two areas of the skull (Fig. 5 C). In contrast to the CFUs (Fig. 1D), at 30 min the number of 1 µm microspheres in the dorsal pachymeninx was higher than in the OB+Skull tissue. The number of 1µm microspheres present in the dorsal pachymeninx was 0.55±0.16% of the number instilled in the nares. This is some 100-fold higher than that of microspheres with diameters of 5 µm (0.0064 ± 0.0004%) and 10 µm (0.0054±0.004%) (Fig. 5 E). Conversely, in the pachymeningeal tissue above the olfactory bulb, no significant differences were found between the three sizes of microspheres (Fig. 5 D). The abundance of 5 and 10 µm microspheres in the pachymeninx of the olfactory bulb, compared to their paucity in the dorsal pachymeninx, with the opposite being true for 1 µm microspheres, shows that the transport of the larger microspheres is hindered. Since the data are limited to one time point, it is not possible to say if the hindrance of the larger microspheres is uniform along the pathway (they travel more slowly), or if it occurs particularly caudad to the meninges of the olfactory bulb.

**Figure 5.**
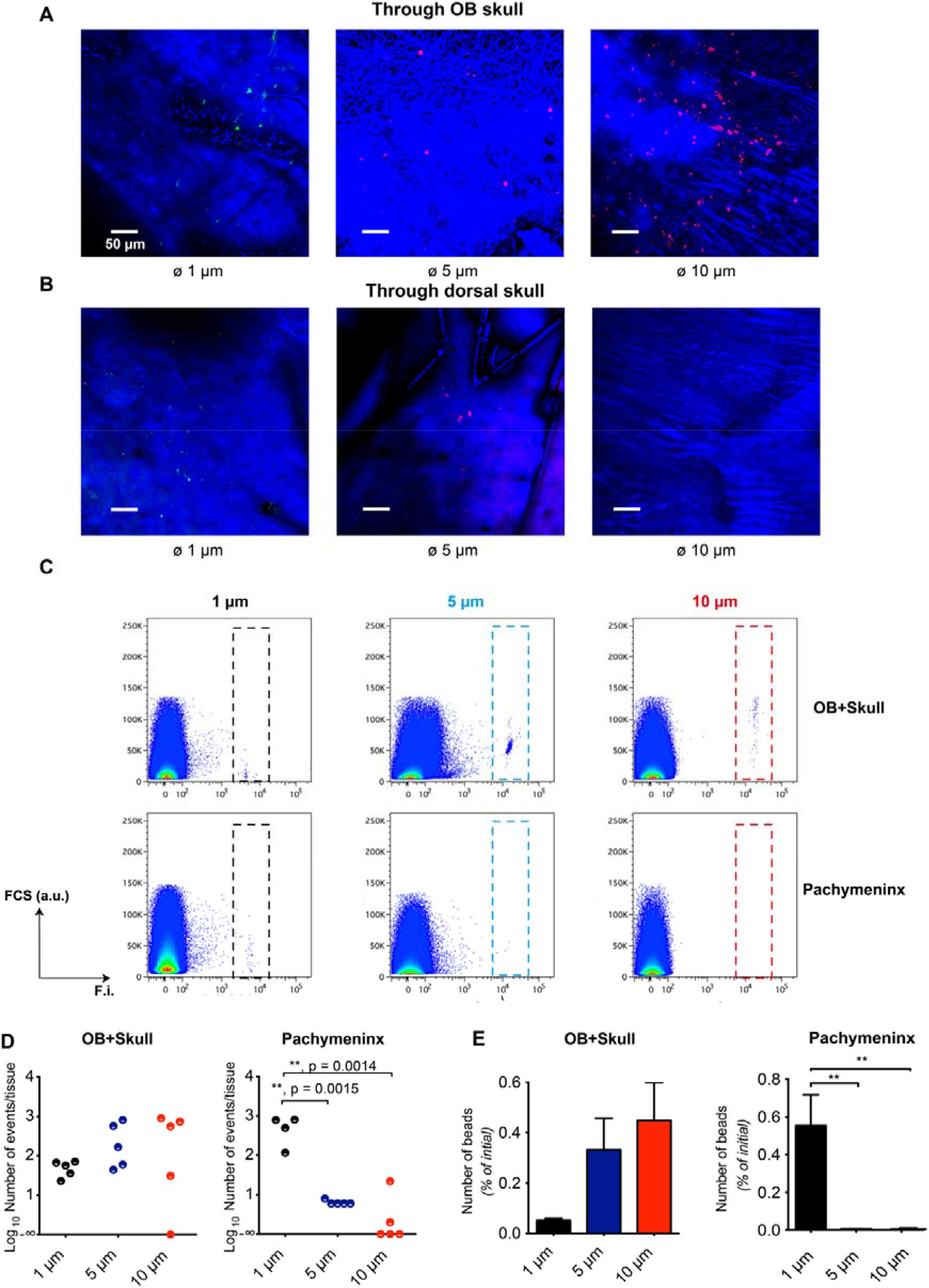
The speed of transit from nose to calvarial meninges is lower for larger microspheres.

To see if Sp and microspheres were passing through the cribriform plate, we removed brain tissue from above the ethmoid bone post-mortem until the tissue was thin enough to allow two-photon imaging of the cribriform plate and its overlying tissue. The foramina were clearly visible (Fig. 6 A), and the overlying collagen of the dura mater appeared also to have holes that could allow passage of olfactory nerve bundles (Fig. 6 B). In other mice, we administered fluorescent Sp or microspheres to the nose, culled the mouse at 15 min. Imaging of the cribriform plate and overlying tissue in a tissue bath showed Sp and microspheres very close to the bone (Fig. 6 C, D). Both Sp and microspheres were present very close to the bone, moving slowly, if at all. This supports the conclusion that Sp and microspheres passed through the cribriform plate and entered the pachymeninx. Some microspheres were in the superfusate, and drifted with its thermal convection (squiggles in Fig. 6 C). This shows that they were not trapped inside cells.

**Figure 6.**
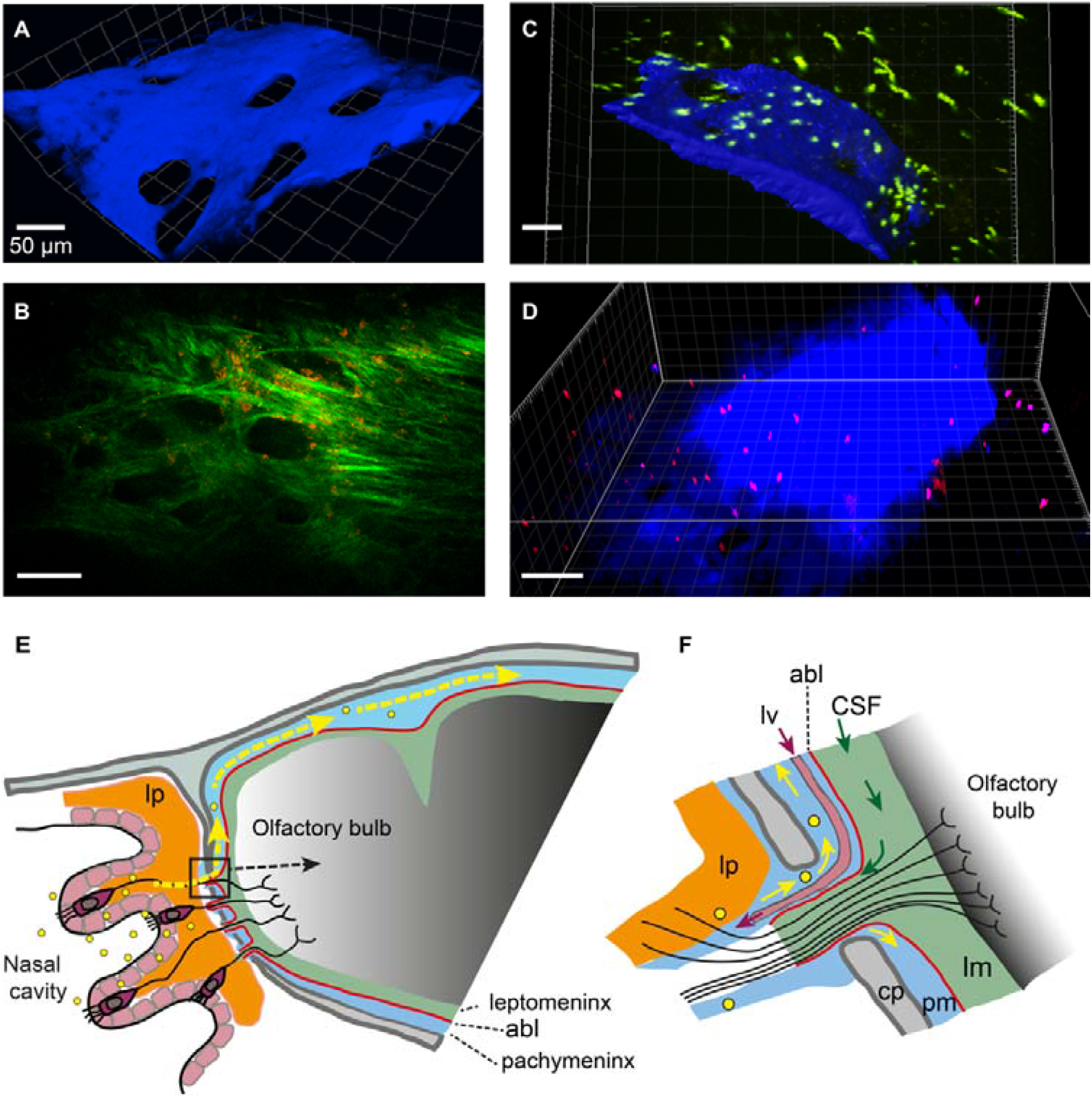
Microspheres and *S. pneumoniae* on the cribriform plate. Two-photon images of the intracranial face of freshly dissected ethmoid bone covered by a layer of olfactory bulb tissue (which gives no signal). (**A**) Excitation at 900 nm gives blue SHG from the cribriform plate. This was a naive LysM-GFP reporter mouse. (**B**) Excitation at 1140 nm gives an image of collagen-like fiber structures, presumably dura mater. This was a CD11c-YFP reporter mouse. YFP is poorly excited at 1140 nm but faint CD11c+ cells can be seen. (**C**,**D**) 3.8 × 10^7^ yellow-green fluorescent polystyrene microspheres of 1 µm diameter size (green) (C) or BacLight™ Red-stained pneumococci (red) (D) were administered intranasally. The mice were culled at 30 min and the intracranial face of the cribriform plate (blue, SHG) imaged with excitation at 840 nm. Baclight™ Red was detected at 571-664 nm. Microspheres and pneumococci are seen close to the bone; some microspheres are drifting in the superfusate. **(E)** Scheme (not to scale) of the anatomy of the pathway (partly hypothetical, sagittal section). Olfactory neurons with their cell bodies in the olfactory epithelium send axons through the foramina of the cribriform plate where they are surrounded by cells which have been described as ‘olfactory ensheathing cells’^11,125-128^ or as forming extensions to some, or all, of the pia, the arachnoid, the dura and the periosteum^27,129-131^. Microspheres and pneumococci (yellow dots) are transported through the cribriform plate and are found in the pachymeninx (yellow arrows) which is separated from the leptomeninx by the arachnoid barrier layer (abl, red line). **(F)** Enlargement of the dashed rectangle in (D). Cerebrospinal fluid (CSF) flows out of the subarachnoid space of the leptomeninx (lm) along extracellular spaces in a bundle of olfactory nerve fibres that traverses a foramen of the cribriform plate (cp). Lymph draining from the pachymeninx flows out ^59,122^ through a lymph vessel (lv), Sp and microspheres are carried into the pachymeninx (pm) along a space adjacent to the lamina propria (lp) (Galeano et al. 2018). The arachnoid barrier layer (abl) is indicated by a red line.

## Discussion

We have shown that in mice infected by horizontal transmission similar to the natural mode of transmission in humans ^43-46^, pneumococci can invade the meninges without detectable bacteraemia (Fig. 1C). Since we were particularly interested in dissemination at short times (minutes) after instillation to the nasopharynx, and also reproducible infections, we had recourse to using nasal instillation, so that the time of infection and the number of Sp applied could be controlled. This procedure is widely used for studying the entry of pathogens into the central nervous system ^7,9,10,33^. We chose a dose of Sp (10^8^ CFU/mouse) that reliably led to dissemination. Although this dose is at the higher end of the range used by most previous studies ^10,46,81^, it should not prevent elucidation of the anatomical pathway at times too early for cell damage to occur; at longer times, there were no signs of major damage to the nasal mucosa, such a bacteraemia. We used a self-limiting model, as used by others, e.g., ^9,10^, in which the administration of pneumococci did not lead to the development of any overt disease symptoms. Hence our results describe the early stages of pneumococcal entry into the central nervous system, upon intranasal administration. These are the early events and entry route that are almost impossible to study in the clinic.

Our key novel finding is the rapid invasion of the pachymeninx by *S. pneumoniae*. We showed that upon intranasal instillation, Sp can reach the pachymeninx very rapidly, within 2 min. We have found no previous studies that have examined the dorsal meninges earlier than one hour after intranasal administration of material of any kind. At one hour, Clark ^82^ found Prussian blue in the pachymeninx, and Galeano et al. ^27^ found stem cells at 2h, as we found for Sp and microspheres (Figs 1D, 2,5). The prime evidence that the Sp were in the pachymeninx, rather than the leptomeninx, is the proximity of Sp to the skull (Fig. 2A-I) and to LYVE-1+ structures ^59,60^ (Fig. 2J-M). This finding is supported by the absence of viable pneumococci in the CSF at 72h post-infection which suggests that Sp did not breach the arachnoid barrier layer and reach the CSF channels in the leptomeninx ^20,25,74,83^ (Fig. 6 E,F). The absence of viable pneumococci in the CSF, combined with the absence of clinical symptoms, raises questions on the pathological importance of our finding. In the clinic, acute bacterial meningitis is not normally diagnosed if viable bacteria are absent from the CSF ^84,85^. However, a number of reports suggest that CSF-negative cultures do not rule out an intracranial bacterial pathology ^86-88^, nor does the absence of clinical symptoms ^89-92^.

The flow cytometry analysis of Ly6G^+^ cells and the intravital imaging of LysM^+^ cells showed that, upon infection, the number and mean speed of LysM^+^ cells in the dorsal pachymeninx increase up to 10h post-infection, these events being hallmarks of local inflammation and immune cell activation. Although this recruitment of LysM^+^ appears to be too slow to account for the rapid fall in CFUs over the first 30 min in the dorsal skull preparation, after the subsequent rebound, the number of CFUs begins to fall again at less than 5h post-infection, as the LysM^+^ population approaches its maximum. It appears therefore, that the LysM^+^ population increases until the clearance of Sp is established ^68,93^, then begins to fall. Intense immune reactions in the pachymeninx (rather than the leptomeninx) have been much studied for their occurrence in migraine^21^, and also observed in experimental autoimmune encephalitis ^94,95^, trypanosomiasis ^41^ and infection by lymphocytic choriomeningitis virus ^72^. The immune cells may have arrived by extravasation from dural vessels ^41,96^, or from the skull bone marrow (Herisson et al. 2018) by way of the transcalvarial channels that contain veins ^73,97,98^. This recruitment of immune cells failed, however, to clear the bacteria and was followed instead by sustained, albeit decreasing, densities of Sp over days (Fig. S2). It remains to be determined what conditions are permissive to the persistence of Sp within the meninges e.g., T regulatory-mediated mechanisms ^99,100^.

We found that 1 μm microspheres, as well as pneumococci (which have about the same diameter), translocate rapidly from the nasopharynx to the pachymeningeal compartment of the dorsal meninges. In the case of microspheres, this compartment was further distinguished from the subarachnoid space by injecting microspheres of a different colour in the cisterna magna; it is known that from there, material is carried by CSF to the dorsal subarachnoid space ^41,63,64^ (Fig. 3). Further, both microspheres and Sp appear to pass through the cribriform plate (Fig. 6 C, D). Hence, we hypothesize that they translocate along the same pathway.

At least five anatomical routes of transport through the foramina of the cribriform plate ^11^ have been proposed: transport within axons (anterograde along the olfactory axons ^101,102^) or retrograde along the trigeminal axons ^103^; transport within the olfactory nerve ensheathing cells ^8,104-106^, transport along extracellular,’perineural’, spaces of the nerves ^8,101,107-109^, transport in exiting lymph vessels ^59,110^ and transport within or close to the periosteum ^27^. Objects as large as pneumococci or micron-sized microspheres diffuse much too slowly for diffusion to account for the rapid transport observed here, so either convection in a flowing fluid, or some form of active transport is necessary. Axonal transport, typically 0.15 mm/min ^111,112^ would take 33 min for a distance of 5 mm from nasopharynx into the cranial meninges, and is therefore also too slow. Further arguments against such an intracellular route are that the olfactory axons are typically only 0.2 μm in diameter ^113^, much less than the diameter of Sp, and that sialic acid, a component of the extracellular glycocalyx of almost all cells ^114^ promotes translocation of Sp to the olfactory bulb ^9,10^, which suggests that interaction with extracellular structures is important for the translocation. As for convection, a puzzle is that numerous results show an efflux of fluid from cranium to nose, rather than an influx ^115-118^. The major conduits for efflux appear to be the spaces between the ensheathing cells that fasciculate the olfactory nerve ^113,119,120^ and the lymph vessels ^59,110^. The former may drain the subarachnoid space ^82,119,120^ and the latter the pachymeninx ^74,121-123^. Although there are reports of extracellular transport from the nasal mucosa towards the cribriform plate along the olfactory nerve ^8,82,110,124^, a third extracellular route, described by Galeano et al. ^27^, along a space between the lamina propria and the turbinate bone, has the merits that it connects directly to the pachymeninx and has not been reported to carry an efflux. Of the known anatomical routes, this is therefore the most probable for the transport of Sp (Fig. 6F), for 1 µm microspheres and perhaps other particulate matter such as pollutants and drugs ^28^ targeting the central nervous system (CNS). Microspheres of diameters 5 and 10 μm were transported more slowly (Fig. 5), suggesting hindrance by the narrowness of spaces or extracellular matrix.

Our study highlights the anatomical structures and fluid networks connecting the nasal cavity to the central nervous system, and their barrier functions. By establishing that both pneumococci and microspheres translocate in minutes from the nasal cavity to the dorsal pachymeninx of mice, our data show for the first time the existence of a previously unrecognised inward flow of fluid through to the CNS. Should the CSF and/or brain parenchyma be subsequently infected, this would mean that pneumococci (and perhaps any microparticles of similar size) would be capable of crossing the arachnoid barrier membrane. The exact mechanisms remain to be determined. Assuming similarities with animal models, our findings have significant implications for the diagnosis and clinical management of CNS infection in human patients. Further investigations into the nose-to-meninges translocation pathway will provide additional insights into not only the nature and the dynamics of host-pathogen interactions in the CNS, but also the development of novel drug delivery systems to the brain, and the etiology of brain damage caused by air-borne particles such as pollutants.

## Supporting information

Audshasai et al_Supplem_BioRxiv

## Authors contributions

T.A., J.A.C., A.K. and M.Y. designed the study. T.A., J.A.C., S.P., S.K., and M.Y. performed experiments. T.A., J.A.C., A.K, and M.Y. wrote the manuscript, and the other authors contributed to data analysis or writing. All authors read and approved the final version of the manuscript.

## Acknowledgments

We acknowledge funding support from Meningitis Now, the UK Medical Research Council (Programme Grant Number MR/P011284/1) awarded to A.K. and the Mahidol-Liverpool PhD Scholarship awarded to T.A. We also acknowledge Joshua I. Gray for assistance with intravenous injections; David Mason, Jennifer Adcott, Dr Marco Marcello, and Dr James Szczerkowski at the University of Liverpool, Centre for Cell Imaging, for assistance with image acquisition and analysis; Dr Lynn MacLaughlin, Sarah Roper and the technical staff at the Biomedical Services Unit, University of Liverpool; and Colin Hughes and the technical staff at the Central Research Facility, University of Glasgow. The authors declare no conflicts of interest.

## Online supplementary material

Supplementary Fig. S1 **Horizontal transmission of *Streptococcus pneumoniae* in mice without Influenza A virus co-administration**. Pneumococcal CFUs determined in various tissues under the same experimental conditions as Fig. 1C when influenza virus A was not co-administered with pneumococci.

Supplementary Fig. S2 **Long-term monitoring of pneumococcal density after intranasal instillation**. Pneumococcal CFUs determined in various tissues at up to 14 days after intranasal administration with *Streptococcus pneumoniae*.

Supplementary Fig. S3 **Fluorescence microscopy images of *Streptococcus pneumoniae***. Two-photon microscopy image of strain 2 serotype D39-loaded with CFSE using (A), and confocal microscopy image of serotype 1/ST217 pneumococci stained with BacLight™ Red (B).

Supplementary Fig. S4 **Confocal images of mouse whole skull mount with transmitted light**. Confocal images in Fig. 2 panels K and L shown here with transmitted light: Sp-infected mouse (A) and uninfected mouse (B).

Supplementary Fig. S5 **Immune cell FACS gating strategy**. Flow cytometry gating strategies used for LysM^+^, CD11c^+^, and Ly6G^+^ cells.

Supplementary movie 1. **Intravital imaging of an uninfected LysM-eGFP mouse**. Two-photon imaging through thinned skull, under the same experimental conditions as Fig. 4D.

Supplementary movie 2. **Intravital imaging of an Sp-infected LysM-eGFP mouse**. Two-photon imaging through thinned skull at 10h post-infection, under the same experimental conditions as Fig. 4C.

Supplementary movie 3. **Intravital imaging of a Sp-infected CD11c-eYFP mouse**. Two-photon imaging through thinned skull at 3.5h post-infection, excitation at 960 nm with x20, N.A. 1.0 water immersion objective. Detector channels were set at <490 nm and 570 nm for second harmonic generation and CD11c, respectively. Z-stacks = 24 µm deep and total acquisition duration = 27 min.

## Abbreviations

BHI: Brain Heart Infusion
CSF: Cerebrospinal Fluid
CSFE: CarboxyFluorescein Succinimidyl Ester
CNS: Central Nervous System
DC: Dendritic Cell
CFU: Colony Forming Unit
GFP: Green Fluorescent Protein
FCS: Fetal Calf Serum
IAV: Influenza A Virus
LSM: Laser Scanning Microscope
N.A.: Numerical Aperture
PBS: phosphate buffered saline
PFA: paraformaldehyde
SHG: Second Harmonic Generation
Sp: *Streptococcus pneumoniae*

