## Supplementary material for "*Streptococcus pneumoniae* rapidly translocates from the nasopharynx through the cribriform plate to invade and inflame the dura": Audshasai et al_Supplem_BioRxiv

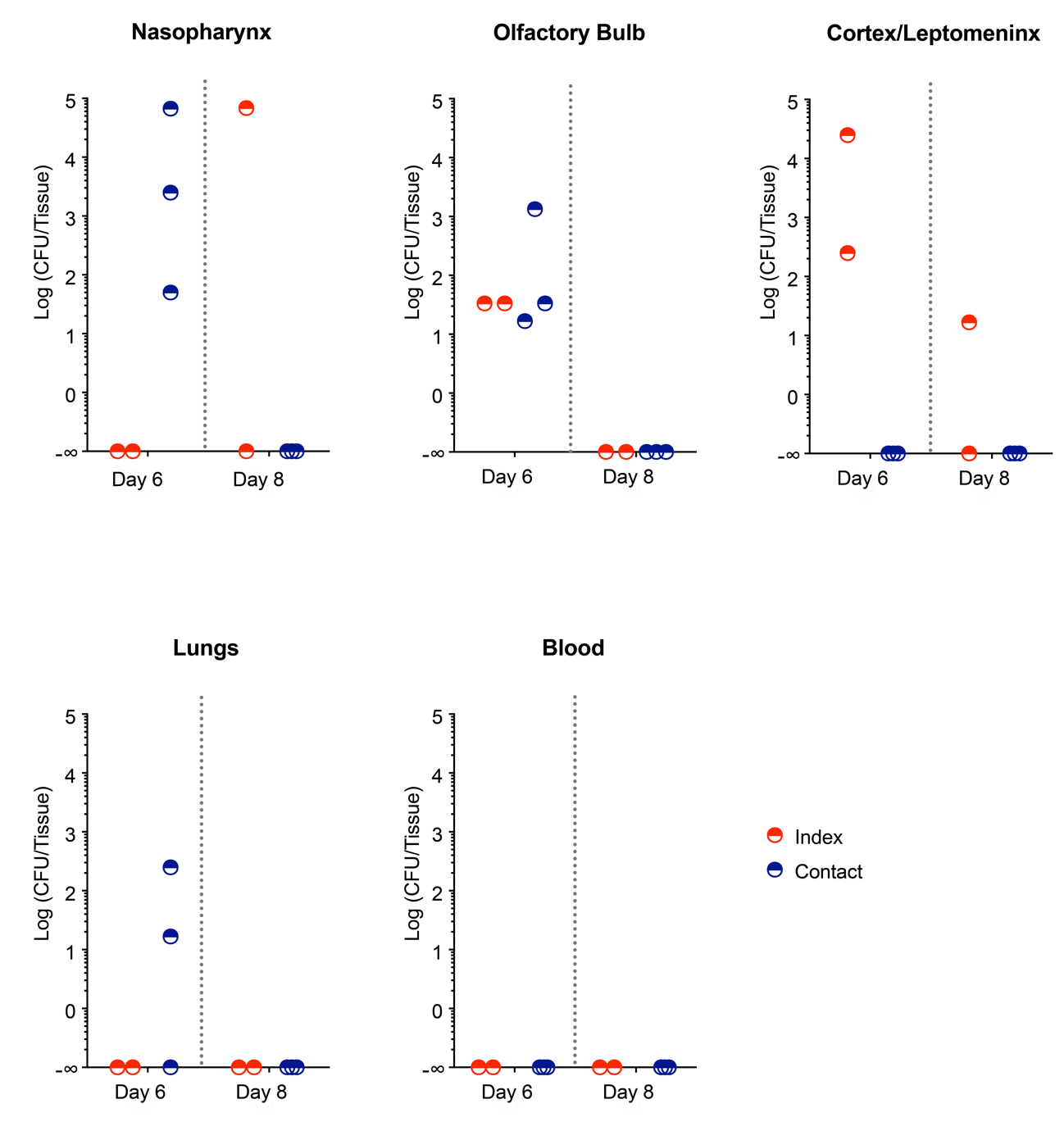

**Supplementary Figure 1.** Horizontal transmission experiments when mice were intranasally administered with pneumococci but without influenza virus A on day 3. Mice were housed in groups of 5 mice per cage. On day 0, two index mice per cage (red symbols) were intranasally administered a 10 µL/mouse inoculum containing approximately 10^5^ CFUs of *S. pneumoniae* serotype-1/ST217 and were returned to the cage with their three contact littermates. On day 6, one group of mice (i.e., 2 index + 3 contact) were killed by asphyxia using CO_2._ Various tissues were examined for CFUs. The procedure was repeated using a second group of mice killed on day 8. Some horizontal transmission was seen on day 6 but not on day 8.

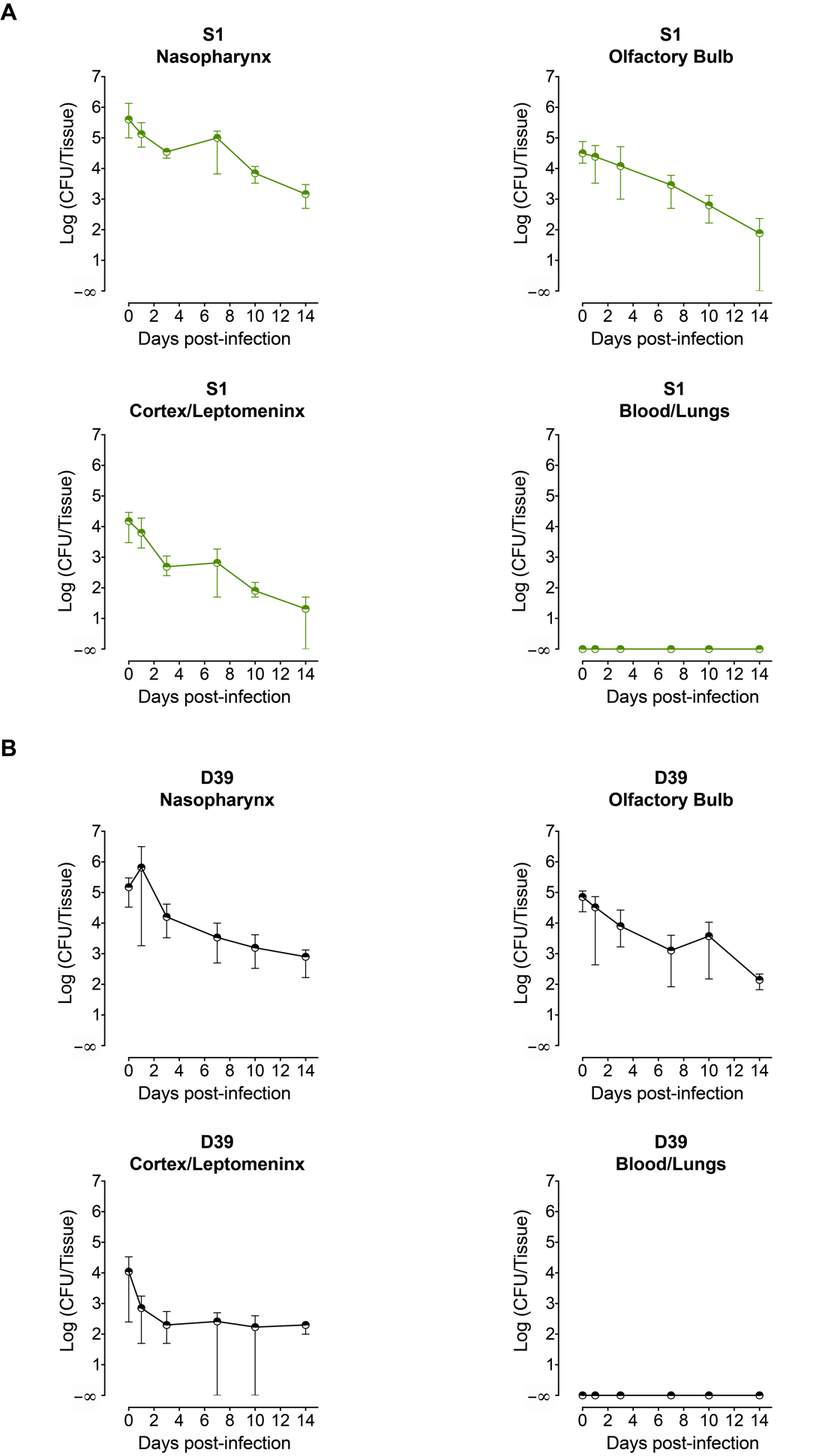

**Supplementary Figure 2.** Pneumococci serotype 1 (S1, sequence type 217) (**A**) or serotype-2*,* strain D39 (NCTC 7466) (**B**) were intranasally administered to mice at a volume of 10 µl/mouse inoculum containing approximately 10^8^ CFUs. CFUs were determined at 15 min, 1 day and subsequently to 14 days post-infection in various tissues. Each symbol represents one mouse, five mice per time. Between day 0 and day 14, an approximate 3-log decline was observed in the CFU counts in all of the collected tissues. S1-infected mice started showing clearance at day 14 in the olfactory bulb (2/5 mice) and in the brain/leptomeninx (3/5 mice). No viable pneumococci were detected in the blood or lung. Note that because no measurements were made between 15 min and 24h any bottleneck at about 1 hour, as in Fig. 1D, would have been missed.

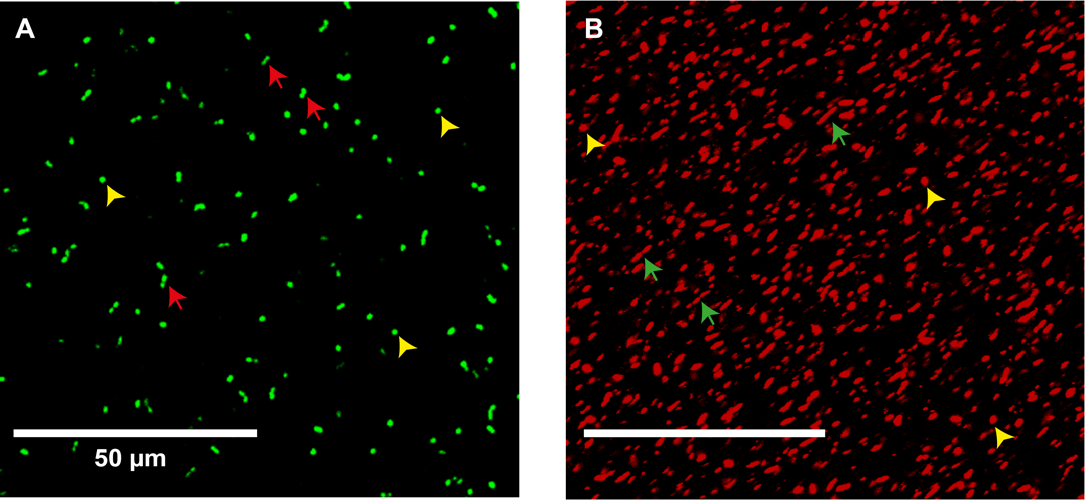

**Supplementary Figure 3.** Two-photon fluorescence image of *S. pneumoniae* serotype 1, sequence type 217 loaded with CFSE for 45 min. The fluorescent *Sp* are bright and clearly distinguishable in either monococci (yellow arrow), diplococci (red arrow) (**A**), Confocal fluorescene image of *S. pneumoniae* Strain , sequence type 217 loaded with Red-fluorescent Baclight^®^ for 15 min. The fluorescent *Sp* are bright and clearly distinguishable in either monococci (yellow arrow), diplococci (green arrow) (**B**).

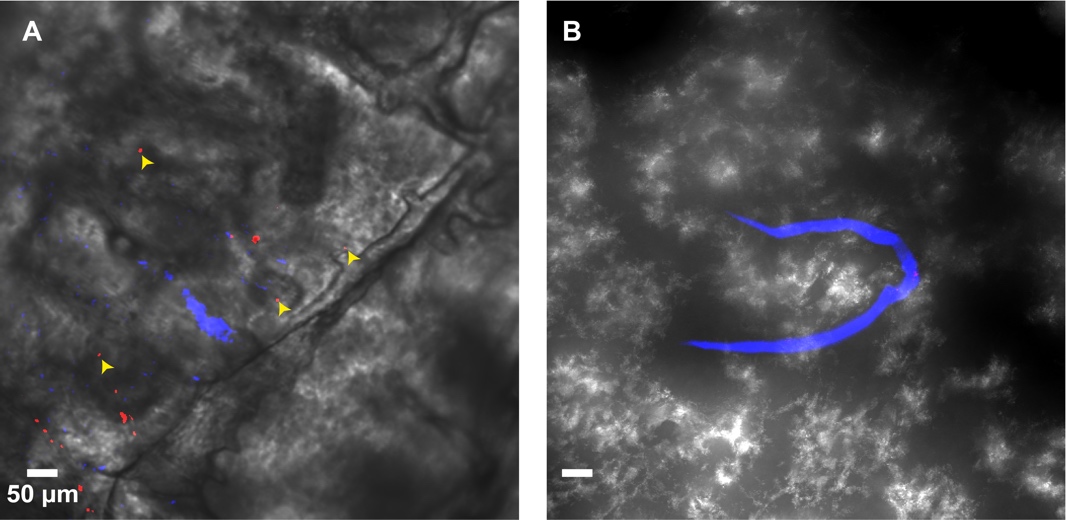

**Supplementary Figure 4.** *S. pneumoniae* stained with BacLight™Red were instilled in the nose. At 15 min post-administration, mice were perfused transcardially with PBS followed by fixing solution (4% PFA). Dorsal skull mounts were stained with anti-LYVE1 antibody and imaged on the skull bone-oriented surface with excitation at 561 nm. A representative Maximum intensity Z- projection of the images of the skull whole mount is shown for the Sp-infected (A) and uninfected (B) mouse with transmitted light included. (Z=239.95 µm).

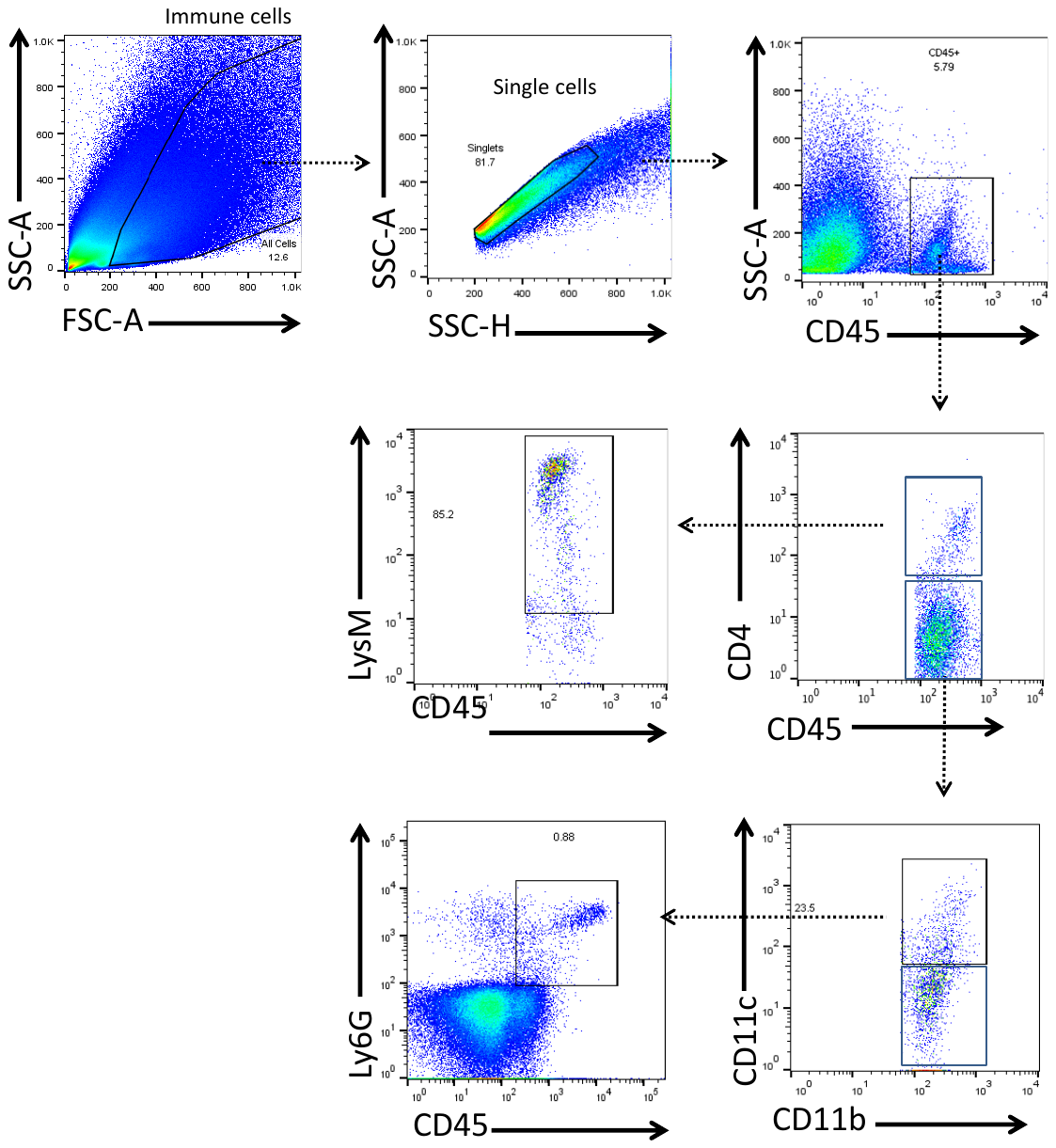

**Supplementary Figure 5**. Flow cytometric discrimination of Ly6G^+^, LysM^+^ and CD11c^+^ cells by surface marker expression. Representative FACS plots, which detail our 6-color flow cytometry gating strategy of single cell suspensions obtained from the mouse pachymeningeal tissues from an Sp-infected mouse at 10 hours post-infection. Neutrophils were defined as CD45^+^ CD4^neg^ CD11b^neg/dim^ Ly6G^+^, myelomonocytic cells were defined as CD45^+^ CD4^neg^ CD11b^neg/dim^ LysM^+^. Dendritic cells were defined as CD45^+^ CD4^neg^ CD11b^high^ CD11c^+^.

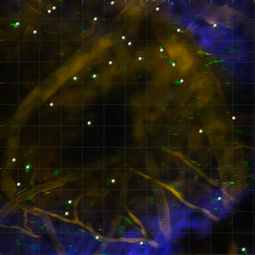
**Supplementary movie 1**. Two-photon intravital imaging of a non-infected LySM-eGFP reporter mouse through thinned skull. LysM+ cells (green) are scarce and present very little movement. Blood microvessels (orange) were stained by intravenous administration of 70 kD dextran-rhodamine. The second harmonic generation (SHG, blue) shows that the LysM+ cells and microvessels are located underneath and very close to the skull at a depth compatible with that of the pachymeninx. Z-projection of stacks 30 µm deep, time series for 32 min. Image field is 424 µm x 424 µm.

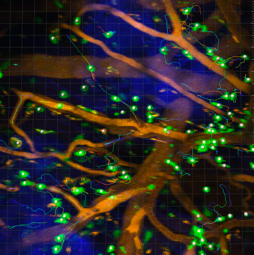
**Supplementary movie 2**. Two-photon intravital imaging of an eGFP-LySM reporter mouse through thinned skull at 10 hours following the intranasal administration of Sp. The settings used for image acquisition and image analysis were identical to those used in the supplementary movie 1. A significantly denser population of LysM+ cells (green) with rapid movements were observed compared to the non-infected mouse. Z-projection of stacks 23 µm deep, time series for 15 min. Image field is 424 µm x 424 µm.

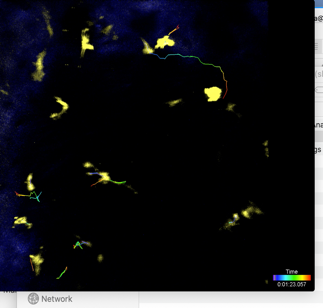
**Supplementary movie 3**. **Supplementary movie 3**. Two-photon intravital imaging of a CD11c-eYFP+ reporter mouse through thinned skull at 3.5 hours following intranasal administration of Sp. CD11c+ cells (yellow) have a morphology characteristics of dendritic cells, with extending and retracting processes but very little displacement. The SHG (blue) shows that the CD11c+ cells are also located underneath and very close to the skull. Z-projection of stacks 24 µm deep, time series for 27 min. Image field is 425 µm x 425 µm.
